# Label-free identification and classification of leukocyte subpopulations in peripheral blood smears by optical photothermal infrared spectroscopy

**DOI:** 10.64898/2026.10.14.759126

**Authors:** Shravan Raghunathan, Susann Piehler, Cornelia Reuter, Michael Kiehntopf, Jürgen Popp, Christoph Krafft

## Abstract

The differential count of leukocyte subpopulations from peripheral blood smears remains a cornerstone of clinical hematology but relies on staining protocols or flow cytometry, that introduce variability and obscure native biochemistry. This study presents label-free workflows that distinguish leukocytes from erythrocytes in full blood smears and classifies the leukocyte subtypes as granulocytes, lymphocytes and monocytes using their single cell optical photothermal infrared (OPTIR) spectra. The first approach uses discrete wavenumber images using two key bands at 1546 and 2922 cm^-1^ to screen leukocytes. The second approach using intrinsic fluorescence imaging of leukocytes offers approximately 1000 times faster screening. Multiple OPTIR spectra are acquired per leukocyte followed by post-acquisition Giemsa staining for cytological cell verification. Full blood smears were prepared on calcium fluoride and standard glass slides to compare substrate effects, and OPTIR spectra were collected with 532 and 785 nm probe lasers to assess probe wavelengths effects. Principal component–linear discriminant analysis separated granulocytes from mononuclear subtypes under all substrate–probe laser combinations tested on the basis of lipid, nucleic-acid and protein bands in OPTIR spectra. Lymphocytes and monocytes were better distinguished on calcium fluoride than on glass. The OPTIR spectra stayed stable over 14 days of storage on both substrates, whereas the intrinsic fluorescence used for cell localization was weak in fresh smears and increased with storage. The non-destructive, fixation-free measurement preserves native protein and lipid composition and allows subsequent cytological staining, supporting OPTIR on routine glass slides for future label-free hematological screening.

## Introduction

The immune system serves as the body’s primary defense against infections and disease, orchestrating its response largely through the activation of leukocytes, also known as white blood cells (WBCs). In clinical diagnostics, the evaluation of a peripheral blood smear, in which leukocytes are identified against a background of erythrocytes (red blood cells, RBCs), remains one of the most widely used hematological procedures.^1,2^ In healthy adults, the three main leukocyte subtypes are neutrophils (35-75%), lymphocytes (18-50%), and monocytes (<12%). Neutrophils are granulocytes, 10-14 µm in diameter with a multilobed nucleus and cytoplasmic granules containing oxidative metabolites and digestive enzymes; their primary functions are phagocytosis and digestion of pathogens.^3,4^ Eosinophils and basophils are less abundant granulocyte populations, typically accounting for approximately 1-5 and 0.5-1% of circulating leukocytes, respectively. Lymphocytes (T cells, B cells, and natural killer cells) are the smallest at 10-12 µm, with a large near-circular nucleus and minimal cytoplasm. Monocytes are larger (15-22 µm) but share a similar mononuclear morphology. Lymphocytes and monocytes form the agranulocyte group. Changes in leukocyte number, morphology, biochemical state, and subtype distribution are associated with infections and other pathological conditions.^5^

The clinical differential count relies on Romanowsky-family stains followed by manual or automated morphological assessment,^1,6^ or by flow cytometry exploiting forward/side scatter and, increasingly, endogenous fluorescence.^7,8^ Although accurate, these approaches either depend on staining variability or report on size and granularity rather than chemical composition. Although fluorescence, traditional flow cytometry, and evaluation of stained blood smear images can provide accurate leukocyte subpopulations, challenges remain regarding specificity and sensitivity to infections, and staining procedures are themselves invasive and can introduce variability in leukocyte labeling and analysis.^9^ A complete understanding of the host immune response, which alters both the number and molecular composition of leukocytes, requires biochemical information at the single cell level. Vibrational spectroscopy offers a label-free, non-destructive, and chemically specific route to this information. Raman spectroscopy has been applied to support high throughput, label free cellomics,^10^ to differentiate fungal and bacterial infections of neutrophils,^11^ to distinguish normal from leukemic hematopoietic cells^12^ and to discriminate neutrophils and lymphocytes.^13^ Infrared (IR) spectroscopy has likewise been used to identify blood and leukemic biomarkers,^14,15^ to discriminate normal human leukocytes from BCR/ABL positive leukemic cells by unsupervised analysis^16^ and to characterize mononuclear leukocytes in peripheral blood from Alzheimer’s disease patients.^17^

Despite these advances, a label-free workflow that integrates leukocyte morphology with chemical bond-specific information at clinically relevant throughput is still lacking. A supervised spectroscopic approach with dynamic leukocyte labeling could help address this gap at the methodological level as a step toward a novel blood smear evaluation. One possible solution is optical photothermal infrared spectroscopy (OPTIR) with optional fluorescence guidance.^18,19^ OPTIR is an indirect IR approach that combines IR excitation by a tunable pulsed quantum cascade laser (QCL) and registration of the resultant photothermal response of the sample due to absorption, by either optical detection of sample expansion^20^ and/or refractive index modulation.^21^ Compared to Fourier transform IR spectroscopy or QCL-coupled direct IR spectroscopy, advantages of OPTIR in the context of single cell analysis include sub-micrometer lateral resolution,^22^ absence of anomalous scattering effects, compatibility with various substrate materials in reflection and transmission modes,^23^ high speed imaging at discrete wavenumbers^24^ and simultaneous collection of IR and Raman spectra.^25^ Moreover, the use of glass slides is more attractive for routine clinical applications, due to their higher robustness and much lower cost compared to calcium fluoride (CaF_2_). This was demonstrated to differentiate malignant and non-malignant lung cells using OPTIR.^26^ Previously, OPTIR was applied to screen leukocytes in a blood smear using discrete wavenumber imaging and unsupervised clustering analysis of OPTIR spectra.^27^ Building on this, the present study introduces supervised, qualitative analysis of leukocytes on both CaF_2_ and routine microscope glass slides.

This methodological study applies OPTIR to leukocytes in unstained peripheral blood smears prepared on glass and CaF_2_ substrates. The aim is to integrate single-cell morphology with chemical bond-specific information in a label-free workflow. First, leukocytes are located in discrete wavenumber OPTIR images. Second, leukocytes are dynamically labelled by their endogenous fluorescence using an enhanced green fluorescent protein (EGFP) filter. Subtype assignments such as granulocyte, lymphocyte, or monocyte are made from nuclear morphology. Single-cell OPTIR spectra are then acquired and analyzed by supervised principal component-linear discriminant analysis (PCA-LDA) of mean spectra. The effects of sample storage and the use of different OPTIR probe wavelengths are also discussed. The study presents a methodological perspective on the use of OPTIR spectroscopy and imaging for cell screening, as a proof-of-concept relevant to future diagnostics of infections and other diseases.

## Experimental

### Sample collection and smear preparation

Blood samples from healthy blood donors were obtained from the Institute of Clinical Transfusion Medicine Jena of the University Hospital Jena after informed consent was given. The study was approved by the local ethics committee (Reg.: 2022-2812-Material). Blood (2.6 ml) was into EDTA tubes (K3EEDTA, plasma-S-Monovettes, 04.1931.001, Sarstedt, Germany) and gently mixed by inverting the tubes immediately after blood collection according to the standard protocols.

### Full blood smears

A drop of EDTA-blood was placed with a pipette onto a microscope glass slide (Thermo Fisher Scientific, Schwerte, Germany) or a CaF_2_ slide (Crystal GmbH, Berlin, Germany) near one end. A cover glass (VWR, Darmstadt, Germany) was placed at approximately 45° on the slide, allowing the drop to spread, and pushed forward in a quick motion to produce a thin film. The smear was air-dried at room temperature.

### Giemsa staining of full blood smears

Air-dried full blood smears were stained following OPTIR measurements according to the Quick-Stain method (Procedure No. GS-10 Sigma/Merck) using Giemsa stain Sigma-Aldrich/Merck KGaA, Germany). Briefly, blood smears were placed in undiluted Giemsa stain for 1–2 min, transferred to deionized water for 2–4 min, rinsed with deionized water, and air-dried afterwards at room temperature.

### OPTIR instrumentation

OPTIR measurements were carried out on two complementary platforms from Photothermal Spectroscopy Corporation (Santa Barbara, CA, USA): the Mirage-LS, used for the leukocyte subtype measurements (CaF_2_ and glass substrates), and the Mirage-R, used for measurements with discrete wavenumber prescreening (glass only). Both instruments share the same core optical configuration, a four-module pulsed quantum-cascade-laser (QCL) pump (Daylight Solutions, San Diego, CA, USA) operating at a 100 kHz repetition rate and a 1% duty cycle, a 40× Cassegrain objective in reflection mode, a detector gain of 1×, a spectral point density of 2 cm^−1^ and a corresponding spectral resolution of 6.6 cm^−1^.

The two platforms differ in three aspects relevant to the present work. First, the pump module on the Mirage-LS (MIRcat QT™) covers 973–1855, 1980–2390 and 2695–2999 cm^−1^ with a 100 ns pulse width, whereas the Mirage-R is equipped with an upgraded module (MIRcat 2400) spanning 934–1800 and 2678–2992 cm^−1^ with a 980 ns pulse width. Second, the Mirage-LS provides two probe lasers (532 and 785 nm) while the Mirage-R provides only the 785 nm probe. Third, the Mirage-LS is additionally equipped with widefield-fluorescence optics (50× objective, EGFP filter set; Green/AlexaFluor 488, Chroma Technology Corporation, Bellows Falls, VT, USA), which were used for cell identification. Across all OPTIR acquisitions the Z-position was determined by the instrument’s autofocus routine, which monitors fluctuations in the OPTIR signal during axial scanning and anchors the focal plane to the substrate–cell interface rather than the cell apex, mitigating reflection-mode artefacts associated with variable cell heights. Reference images of the Giemsa-stained leukocytes were collected on a Keyence BZ-X810 microscope using a 40× objective at a resolution of 0.04 µm per pixel.

### OPTIR spectral and image acquisition

For the peripheral blood smears using the Mirage-LS, five OPTIR spectra were acquired per individual leukocyte with two co-added scans per spectrum (three from the nuclear region and two from the cytoplasm wherever possible) and subsequently averaged to capture the biochemical contributions of the whole cell. Spectra were recorded in the high-SNR mode at a sweep speed of 200 cm^−1^/s. The pump laser operated at 61% (2.8 mW) of its average power, and the probe laser at 10% (8.7 mW) for the 532 nm line and 35% (7.8 mW) for the 785 nm line. Wide-field fluorescence images were collected with the EGFP filter set at 100 ms exposure. OPTIR spectra were acquired from leukocytes under three configurations: probed at 532 nm on CaF_2_ (50 granulocytes, 30 lymphocytes, 20 monocytes, from single blood smear) and on glass (100, 60 and 40 cells from two blood smears); probed at 785 nm on glass (260, 203 and 73 cells, from seven donors). A full breakdown of donors, smears, and per-subtype cell counts per configuration is given in Table **S1**. Ten-fold cross-validation was performed at the cell level, with folds drawn within each smear of each donor. To test the effect of storage time, smears were kept at room temperature in the dark and measured within 48 hours, after 7 days and after 14 days. At each time point the same set of about ten randomly selected leukocytes was relocated, and re-measured on CaF_2_ and glass, with the 532 nm probe using the settings mentioned above.

For the peripheral blood smear dataset with the Mirage-R, single-cell OPTIR spectra were collected as four co-added scans at 47% pump power (0.6 mW) and 52% probe power (15 mW) using the 785 nm probe. In addition to single-cell spectra, discrete-wavenumber OPTIR images of 1200×1200 µm^2^ fields of view were collected at 1546 cm^−1^ and 2922 cm^-1^ at a step size of 2 µm/point and a scan speed of 2000 µm/s.

### Spectral pre-processing

Pre-processing was performed in a sequential workflow shared between the two datasets, with sample-specific adaptations described below. The common sequence comprised high-frequency noise reduction with a Savitzky–Golay (SG) filter (window 11, polynomial order 2, derivative 0) and baseline correction using the Statistics-sensitive Non-linear Iterative Peak-clipping (SNIP) algorithm with 40 iterations. Pre-processing of the leukocyte-classification (Mirage-LS) dataset was performed in RAMANMETRIX^28^ (v0.6.8); pre-processing of the Mirage-R dataset was performed in Python 3.12 (NumPy, SciPy, scikit-learn) using the same parameter values for SG and SNIP.

For the dataset acquired using Mirage-LS, pre-processed spectra underwent vector (L2) normalization; the spectral region between 973 and 1311 cm^−1^ was excluded due to strong glass-background contribution. For visualization, the second derivative was calculated using the *spectramap*^29^ Python package with SG parameters (11, 2, 2) and multiplied by −1 (polarity-reversed convention) so that genuine absorbance bands appear as positive bands, consistent with the sign convention used throughout this work.

### PCA–LDA classification of leukocytes

The normalized leukocyte spectra were subjected to a supervised principal component– linear discriminant analysis (PCA–LDA) model. The dimensionality of the spectral data was first reduced using the first seven principal components for cells on glass and the first five for cells on CaF_2_, capturing more than 95% of the variance in each case. Linear discriminant analysis was then applied to identify latent variables that best discriminate between the predefined classes (G—granulocytes, L—lymphocytes and M—monocytes). The PCA–LDA model was evaluated by 10-fold cross-validation at the cell level, with folds drawn within in each smear of each donor. Leave-one-donor-out validation was not applied, because the CaF_2_/532 nm (1 donor) and glass/532 nm (2 donors) configurations cannot support it. In the 7-donor glass/785 nm set, the per-donor monocyte count of just ∼10 cells, corresponding to the physiological <5% abundance of this subtype, is too low for a stable estimate. The reported accuracies therefore quantify within-donor spectral learnability rather than independent-donor generalization.

## Results

### OPTIR spectra and images of blood smears on glass

Fig. 1 shows the mean OPTIR spectra acquired with the Mirage-R (785 nm probe laser) of erythrocytes and visually detected leukocytes in a blood smear on a glass slide, together with the negative second derivative spectra. Main IR bands are assigned to the protein amide I at 1658 cm^-1^ and amide II at 1542 cm^-1^. The amide I band is slightly narrower and the amide II band slightly more intense for erythrocytes compared to leukocytes. Both the mean and negative second derivative spectra were normalized at the amide I band (1658 cm^-1^). Further bands at 1452 cm^- 1^ from symmetric bending modes of CH_2/3_ groups, and at 1394 cm^-1^ from stretching modes of COO^−^ groups are broad and virtually unchanged for leukocytes and erythrocytes. The CH_2_ stretching vibrations at 2854 and 2926 cm^-1^ are elevated for leukocytes and can be considered as marker bands whereas the CH_3_ stretching vibrations, at 2874 and 2956 cm^-1^ have similar intensities. The expected nucleic acid bands of the leukocyte near 1080 and 1230 cm^-1^ overlap with broad spectral contributions of glass with large standard deviations. The second derivative spectra of the leukocyte and erythrocyte further resolve overlapping IR bands. They are presented on a negative scale to maintain polarity of band positions with the mean spectra. The broad glass background from 900 to 1200 cm^-1^ is effectively suppressed, which enables to resolve the nucleic acid band of leukocytes at 1080 cm^-1^. The irregularities around 1210 cm^-1^ are caused by low IR intensities at a QCL chip transition which gives reduced signal to noise ratios.

**Figure 1.**
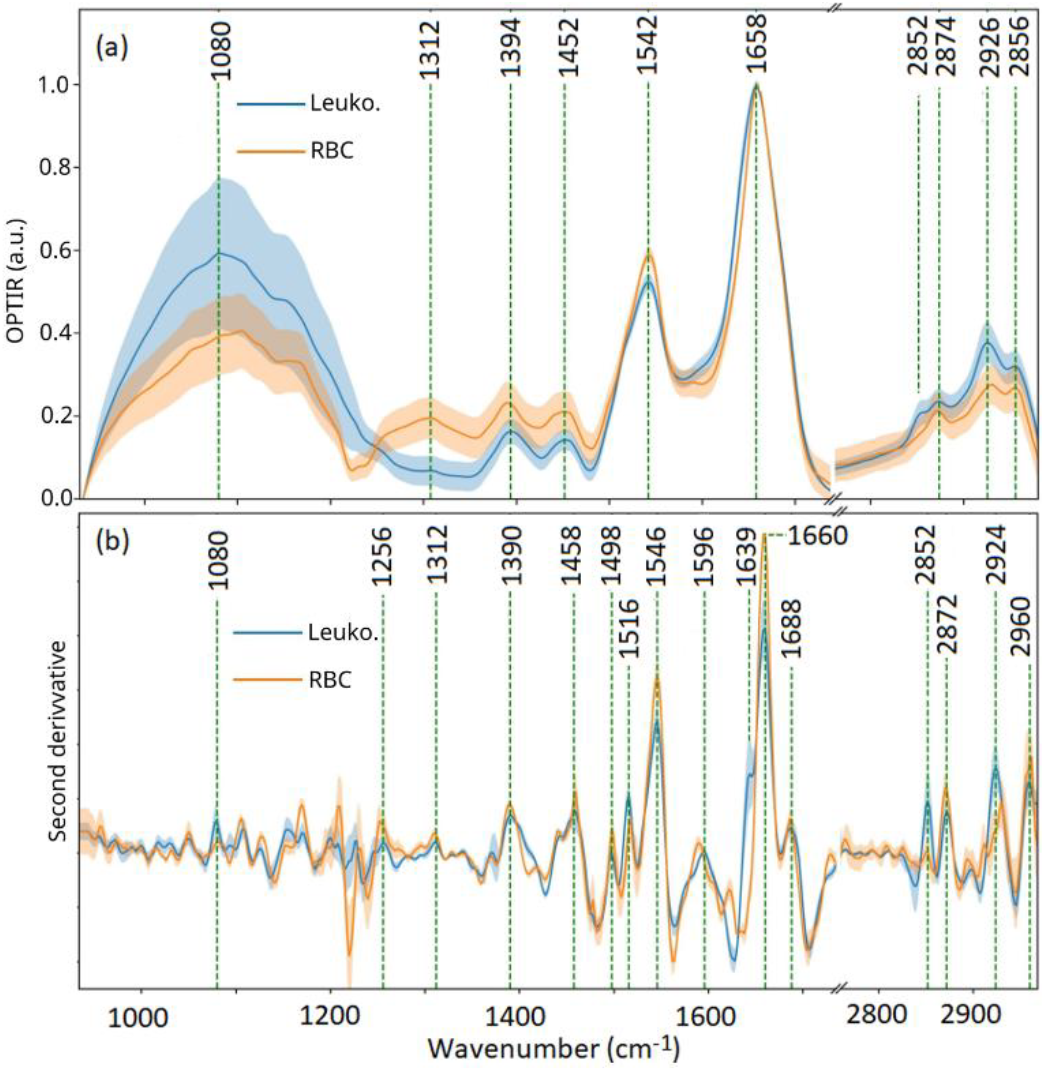
(a) Mean leukocyte (blue) and erythrocyte (orange) OPTIR spectra of blood smears on glass with 785 nm probe laser. (b) Negative second derivative spectra of leukocyte (blue) and erythrocyte (red blood cell, RBC orange). Standard deviations are indicated as shaded regions. Spectrum in (a) and (b) normalized at the amide I band, 1658 cm^-1^.

The smaller width of the amide I band in erythrocytes gives a more intense second derivative band at 1660 cm^-1^ (Fig. 1b). The amide I shoulder in leukocytes is resolved as second derivative band at 1640 cm^-1^, and its larger standard deviation is consistent with different leukocyte subtypes as detailed below in Fig. 5. More sub-bands are resolved at 1256, 1312, 1498, 1516 and 1688 cm^-1^. Key features in second derivatives of leukocytes are more intense bands at 2922 and 2852 cm^-1^ of antisymmetric and symmetric CH_2_ stretching vibrations. All changes for leukocytes are attributed to higher content of lipids and phospholipids in the organelles of cytoplasm and the cell membrane, DNA in cell nuclei, and protein secondary structure variations. Band positions in mean and second derivative spectra are summarized in Table **S1**.

OPTIR images at 1546 cm^-1^ corresponding to the amide II and at 2922 cm^-1^ corresponding to the CH_2_ antisymmetric stretching mode are shown in Fig. 2(b) and (d), respectively. The choice of these OPTIR discrete wavenumbers was based on the observation that erythrocytes show an increased amide II absorption while the leukocytes show an elevated absorption at 2922 cm^-1^ as evident in Fig. 1. A region of interest of size 1200×1200 µm^2^ was chosen for the IR image acquisition from a field of view shown in Fig. 2(a), the step size was set to 2 µm/point and at a speed of 2000 µm/s. The total time taken for the two images was approximately 18 minutes. A properly scaled ratio of the two IR images enables to identify leukocytes in Fig. 2(d). In both images, lower absorptions areas (below∼ 2.5 and 3 mV) were set to zero by setting a thresholding limit. Once the ratio image was evaluated, the ratio in the range of 0 to 1 was masked, which helps in identifying eleven leukocytes as large bright spots in Fig. 2(c).

**Figure 2.**
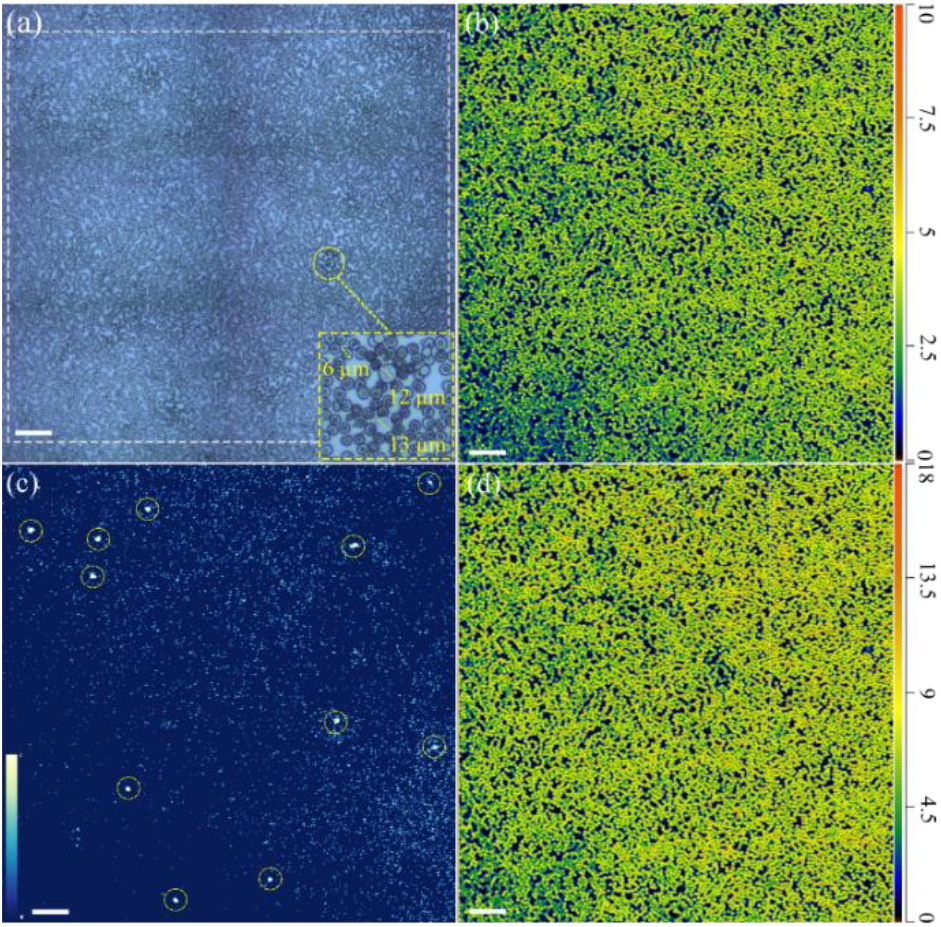
(a) Microscope image of blood smears on glass at 10× magnification. The region of OPTIR image acquisition is highlighted. The Inset shows zoomed-in image of two leukocytes of size = 12 and 13 µm and a single erythrocyte of size = 6 µm. (b) OPTIR image of unstained blood smears acquired at 2922 cm^-1^ (intensity scale in mV). (c) Ratio of 2922 cm^-1^ and 1546 cm^-1^ OPTIR images. The yellow circles point to leukocytes identified. (d) OPTIR image at 1546 cm^-1^ (intensity scale in mV). Scale bar in (a)–(d) = 100 µm.

### Fluorescence and stained images of leukocytes

Although OPTIR images at two discrete wavenumbers enable localization of leukocytes, the speed was not sufficient for high throughput screening. Acquiring two discrete wavenumber OPTIR images over a 1.2×1.2 mm^2^ field of view (FOV) required ca. 18 minutes (1080 s) whereas a wide-field fluorescence image of a comparable FOV can be collected in about less than a second which is more than three orders of magnitude faster. The intrinsic fluorescence of leukocytes with blue and green excitation (450-550 nm) was therefore adopted as a rapid localization option. Fig. 3 shows representative images of the three principal leukocyte subpopulations: (a, a′) granulocytes, (b, b′) lymphocytes and (c, c′) monocytes, captured at two complementary stages of the workflow. The upper panels (a, b, c) were acquired label-free through an EGFP filter and using the 50× objective of the Mirage-LS microscope, prior to staining (fixation), exploiting the intrinsic green fluorescence of intracellular flavins (predominantly flavin adenine dinucleotide, FAD) and, in granulocytes, eosinophilic granule pigments.

**Figure 3.**
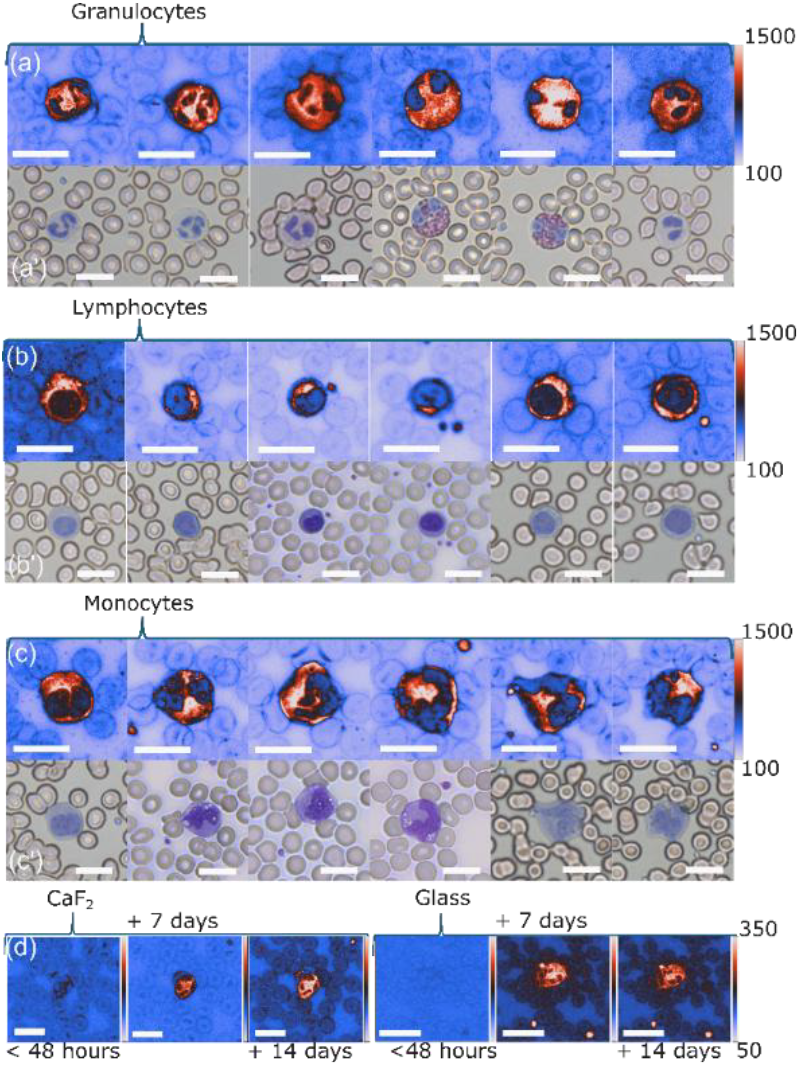
Representative images of single (a) granulocytes, (b) lymphocytes, and (c) monocytes. Top rows (a, b, c): EGFP filter fluorescence; bottom rows (a’, b’, c’): Giemsa-stained brightfield. (d) Leukocyte autofluorescence acquired as function of time. Scale bars in (a - d) = 15 µm.

Critically, this fluorescence-guided step enabled unambiguous targeting of individual cells for OPTIR acquisition without introducing exogenous chromophores that would otherwise contaminate the mid-IR fingerprint region (Fig. **S1**). Panels (a′, b′, c′) show the same fields re-imaged in brightfield mode after post-OPTIR Giemsa staining, which confirmed cell identity based on the following established morphological features: lobulated nuclei with cytoplasmic granularity (granulocytes), a high nucleus-to-cytoplasm ratio with a thin rim of blue-stained cytoplasm (lymphocytes), and eccentric, indented, kidney-shaped nuclei with abundant ground-glass cytoplasm (monocytes). Cells from both the CaF_2_ and glass slides are pooled in the representation. No systematic morphological bias was observed between the two, indicating that the workflow is independent of which one is used.

Another interesting feature of leukocyte autofluorescence is its time dependence. Fig. 3d shows the intrinsic fluorescence of a representative leukocyte on CaF_2_ and on glass, imaged within 48 hours, after 7 days and after 14 days of storage. The fluorescence intensity increased steadily over the 14 days on both substrates, because the smears are dried at room temperature, active cellular metabolism has ceased by the time of the first experiment, and the rise is unlikely to reflect metabolic or redox dynamics. A more plausible origin is slow post-drying photophysical and oxidative aging of endogenous fluorophores, in particular flavins and other green emitting components of the cytoplasm and granules whose emission is known to evolve with progressive oxidation, dehydration and photoproduct accumulation in the fixed or dried preparations.^30,31^ The trend is gradual and reproducible on both substrates, so the brighter signal of aged smears provides clear, label-free contrast for locating cells. For freshly prepared smears, where the fluorescence is still weak, cell localization can instead rely on discrete-wavenumber OPTIR imaging at the amide II (1546 cm^−1^) and CH_2_ (2922 cm^−1^) bands demonstrated above (Fig. 2), or on other contrast methods chosen so as not to alter the native molecular composition probed by OPTIR. The two strategies are complementary and together cover the full storage window.

### OPTIR spectra of leukocytes

Fig. 4 shows the mean and standard deviation of the OPTIR cell spectra obtained from three different leukocyte types: (a) granulocytes, (b) lymphocytes and (c) monocytes. The three different configurations are indicated as cells on glass with 785 nm probe laser (black), on CaF_2_ with 532 nm probe laser (red), and on glass with 532 nm probe laser (blue). The standard deviations in the OPTIR spectra represent cell-to-cell variation after normalization. As observed already in Fig. 1, standard deviations are elevated in OPTIR spectra of cells on glass with 785 nm probe laser. Consistent IR bands are evident at 1657, 1541, 1455, 1391, 1233 and 1081 cm^−1^ in the fingerprint region,^24^ and at 2961, 2925, 2873, and 2855 cm^−1^ in the high-wavenumber region.^32^ The band at 1233 cm^−1^, resolved only on CaF_2_ for all leukocyte subtypes, is due to the antisymmetric stretching mode of PO_2_^−^, found in nucleic acids. The band at 1081 cm^−1^ for the leukocytes on CaF_2_ and glass probed by the 532 nm laser, can be assigned to the symmetric stretching mode of PO_2_^−^. Different from the spectra in Fig. 1, the maximum of the glass band probed by the 785 nm laser is shifted to 1109 cm^−1^. To exclude confounding effects of glass background variations, the spectral range 973-1311 cm^−1^ was not considered for further analysis. In the high-wavenumber region, the symmetric and antisymmetric CH_2_ stretching modes at 2855 and 2925 cm^−1^, the symmetric and antisymmetric CH_3_ stretching modes at 2873 and 2961 cm^−1^ occur at similar positions as in Fig. 1.

**Figure 4.**
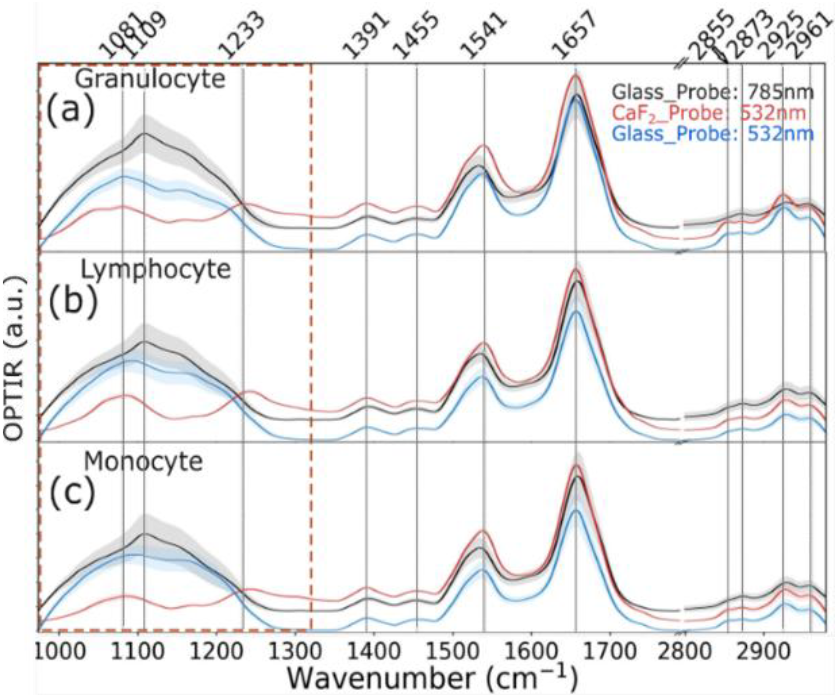
Mean OPTIR spectra of (a) granulocytes, (b) lymphocytes and (c) monocytes acquired under three configurations: on glass with 785 nm probe laser (black), on CaF_2_ with 532 nm probe laser (red), and on glass with 532 nm probe laser (blue). Shaded bands indicate the standard deviation. Red dashed rectangle highlights the region from 973 to 1311 cm^-1^, excluded for subsequent chemometric analyses.

Unlike the intrinsic fluorescence, which increased with storage time (Fig. 3d), the OPTIR spectra of the same leukocytes were stable over 14 days. Mean OPTIR spectra and negative second derivatives are shown collected within 48 hours, after 7 days and after 14 days (Fig. S2). They overlapped within the cell-to-cell standard deviation on both CaF_2_ and glass, with the amide I, amide II and CH-stretching bands unchanged in position and relative intensity. The biochemical contrast used for classification is therefore kept well beyond the immediate evaluation window, so spectra can be acquired on stored smears without loss of subtype information.

### Classification of leukocyte subtypes

To understand whether the three leukocyte subpopulations could be resolved in a supervised low-dimensional space, principal component analysis followed by linear discriminant analysis (PCA-LDA) with 10-fold cross-validation was performed on the mean OPTIR spectra of each leukocyte. The analysis was performed under three configurations: CaF_2_ with 532 nm probe (Fig. 5a-c), glass with 532 nm probe (Fig. 5d-f) and with a 785 nm probe (Fig. 3a-c). For each configuration, Fig. 5a, b and d, e and Fig. S3a, b display the LD1 and LD2 score distributions across granulocytes (grey), lymphocytes (teal) and monocytes (red) as violin plots with embedded boxplots. Fig. 5c, f and Fig. S3c show the corresponding LD1, and LD2 loadings (black) overlaid with the mean negative second-derivative spectra of each subpopulation.

**Figure 5.**
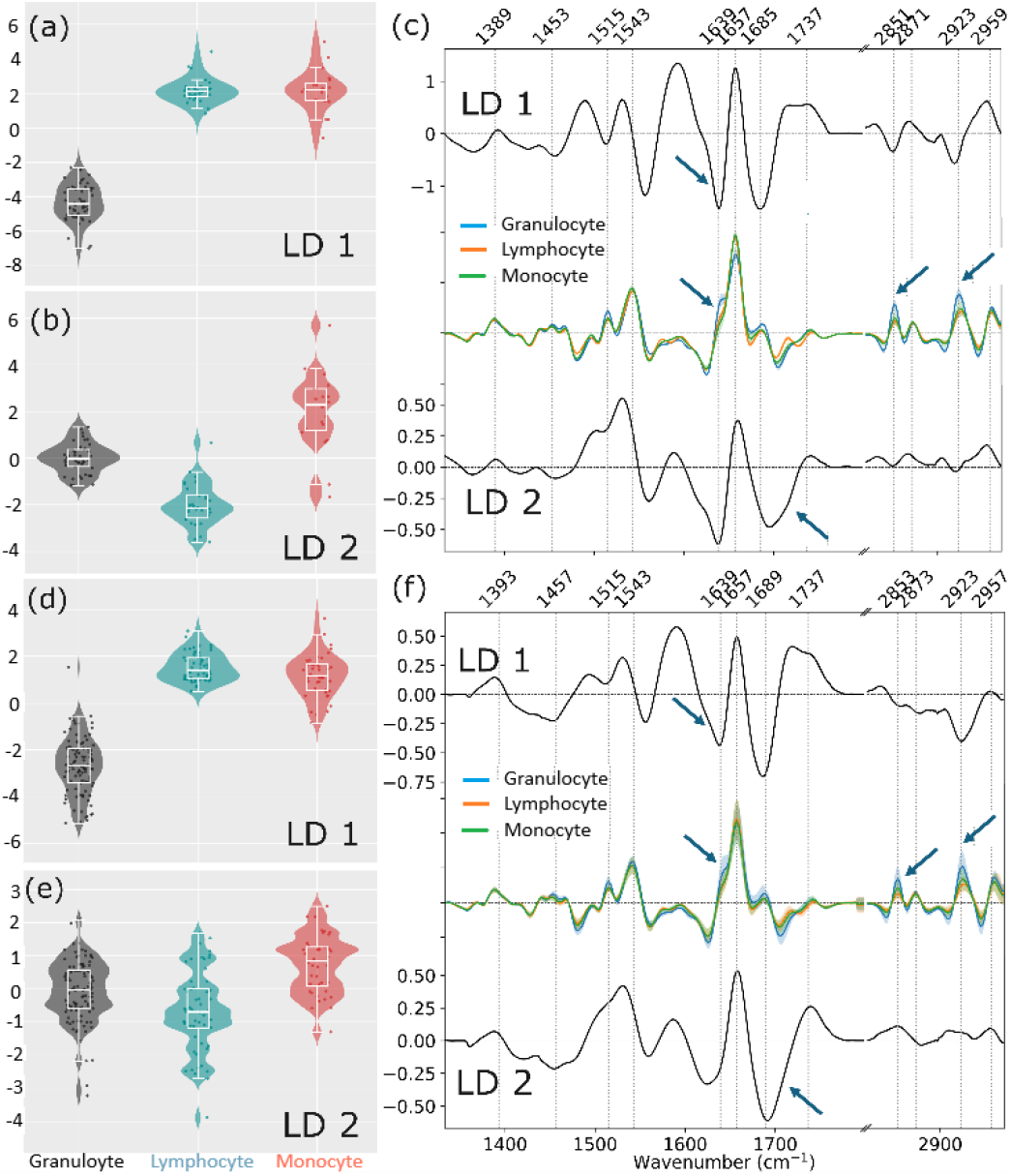
Principal component-linear discriminant analysis (PCA-LDA) analysis of OPTIR spectra from granulocytes (grey), lymphocytes (teal) and monocytes (red). Leukocytes measured on (a–c) CaF_2_ and (d–f) glass with a 532 nm probe respectively. (a, b, d, e) Violin plots of LD1 and LD2 scores across the three subpopulations. (c, f) LD1 and LD2 loadings (black) shown alongside the overlaid mean negative second-derivative spectra of each leukocyte subpopulation for comparison. Arrows highlight the spectral regions driving class separation.

Across all three configurations, LD1 segregated granulocytes from the agranulocyte branch (lymphocytes and monocytes). In the CaF_2_/532 nm configuration, granulocytes occupied a tight negative lobe (median LD1 = −4.41, interquartile range (IQR) 1.53), while lymphocytes (median 2.12, IQR 0.53) and monocytes (median 2.20, IQR 0.96) were displaced to positive scores. The same topology was reproduced on glass at 532 nm (granulocyte median −2.67, lymphocyte 1.41, monocyte 1.16) and 785 nm, confirming that the granulocyte/agranulocyte axis is a substrate- and probe laser-independent signature. Ten-fold cross-validated balanced accuracies of 0.96, 0.80 and 0.63 were obtained for the CaF_2_/532 nm, glass/532 nm and glass/785 nm configurations, respectively. These figures reflect within-donor, cell-level performance. The modest donor pools (1, 2, 7 donors, respectively) do not support a formal donor-level generalization estimate, and in particular the glass/785 nm value should be read as an indication of spectral learnability at that substrate and probe combination rather than a measure of transferability to independent donors.

LD2, in contrast, behaved in a strongly substrate-dependent manner. On CaF_2_/532 nm it resolved the residual lymphocyte-monocyte boundary that LD1 had left ambiguous, with lymphocytes displaced to negative scores (median −2.15, IQR 0.96) and monocytes to clearly positive scores (median +2.29, IQR 1.77) while granulocytes remained centered near zero (median −0.03, IQR 0.73), a three-way separation that accounts for the high balanced accuracy. On glass, however, LD2 collapsed to a much weaker axis, recovering only a marginal monocyte-positive shift (532 nm: granulocytes −0.03, lymphocytes −0.69, monocytes +0.84; 785 nm; all medians within ±0.21 of zero). Taken together, PCA-LDA establishes a robust granulocyte versus agranulocyte axis across all configurations, and a clean lymphocyte-monocyte resolution is realized only on CaF_2_. Granulocytes display pronounced biochemical characteristics including granule-associated lipids, myeloperoxidase, and multilobed, nucleic-acid-rich lobulated nuclei, whereas the two mononuclear subtypes exhibit more closely related biochemical profiles.

The LD loadings (Fig. 5c, f) highlight the spectral basis of the separation, and their coincidence with bands in negative second-derivative spectra confirms that each contribution maps onto a real vibrational mode. Plotting the loadings against the polarity-reversed second derivatives is deliberate, the negative second derivative restores the directionality of absorbance bands while sharpening overlapping bands. The coincident maxima between loadings and spectra directly confirm that the discriminant axes are driven by genuine vibrational features rather than baseline artefacts or spectral noise. The LD1 axis is dominated by features across the amide I envelope, where the loading resolves three closely spaced components (full band assignments compiled in Table S2): 1657 cm^−1^ (α-helix amide I), 1639 and 1689 cm^−1^ (β-sheet amide I). Their opposing signs in the loading indicate that the granulocyte/agranulocyte contrast is not encoded by total protein content but by the relative balance of α-helical versus β-sheet secondary structure. Amide II bands at 1543 and 1515 cm^−1^ also contribute to the discrimination. Further contributions at 1737 cm^−1^ (ester C=O of phospholipids) and across the CH-stretching region at 2853 and 2923 cm^−1^ (ν_s_ and ν_as_ CH_2_, mainly lipids), 2873 and 2957 cm^−1^ (ν_s_ and ν_as_ CH_3_, mainly proteins) further reveal that the granulocyte signature carries a distinctive lipid fingerprint, consistent with the abundance of membrane-rich granules in this lineage. The LD2 loadings recruit a partially overlapping set of bands, but with the amide II shoulder 1515 cm^−1^ and the 1639/1657 cm^−1^ pair carrying the heaviest weight on CaF_2_, in line with LD2’s role in resolving the lymphocyte-monocyte boundary. On glass at 532 nm, the same loadings appear at slightly different amplitudes. The corresponding loadings for the glass/785 nm dataset (Fig. S3c) recruit the same band set with attenuated amplitudes, consistent with the reduced classification performance under this configuration; the underlying biochemical contrast is preserved while spectroscopic sensitivity is diminished at this substrate–probe combination.

## Discussion

Leukocytes and erythrocytes can be distinguished in blood smears on CaF_2_ and glass by discrete wavenumber OPTIR imaging at 1546 and 2922 cm^-1^ which corresponds to the main discriminatory features between these cell classes. The required time of approximately 18 minutes for a 1.2×1.2 mm^2^ FOV is not fast enough to count leukocytes in the monolayer zone in a statistically significant number. A commercial OPTIR version has been introduced with a laser-scanning microscope offering a 30-times speed advantage, and an experimental version reported an imaging line rate of more than 2 kHz in FOVs >200×200 µm^2^.^33^ Wide-field OPTIR, also called mid-infrared photothermal (MIP), instruments enable to collect discrete wavenumber IR images within seconds. FOVs of >1 mm diameter have recently been reported for wide-field OPTIR imaging using computational FOV extension^34^ and an IR laser with pulse energies above 100 µJ.^35^ Fluorescence imaging offers comparable FOVs at similar collection time in the sub-second range using state-of-the-art microscopes. The integrated fluorescence option of the Mirage-LS OPTIR instrument was applied to identify leukocytes in blood smears which constituted a convenient and rapid way to localize them and collect OPTIR spectra. The fluorescence images were used to assign leukocytes to the subtypes granulocytes, lymphocytes and monocytes. This information enabled to train the supervised classification model PCA-LDA which was validated by an assessment of the same cells after Giemsa staining.

PCA-LDA score plots of the OPTIR spectra separate the major three leukocyte subtypes along two axes of molecular contrast. The first and dominant axis, captured by LD1, separates granulocytes from agranulocytes based on protein and lipid features. Protein side chain vibrations at 1393 cm^−1^ are common to all three cell types.^36^ LD1 loadings of granulocytes exhibit negative CH_2_ scissoring at 1457 cm^−1^, CH_2_ stretching at 2851 and 2923 cm^−1^, and amide I sub-bands at 1639 and 1685 cm^-1^ that are consistent changes in the overlaid second derivatives. These features can be assigned to cytoplasmic granules, which are rich in phospholipid membranes and neutral lipids.^36,37^ The second axis, captured by LD2, discriminates lymphocytes from monocytes. The LD2 loadings of lymphocytes show spectral changes in the amide II and I bands with different relative amplitudes and a shoulder near 1690 cm^-1^ pointing to nucleic acid base C=O contributions.^36,38^ This spectral profile is consistent with their high nucleus to cytoplasm ratio and the predominance of proteins over lipids in their cytoplasm.^37^ Monocytes occupy an intermediate position between lymphocytes and granulocytes in the LD score space, reflecting their mononuclear morphology combined with moderate cytoplasmic complexity and accounting for the partial overlap observed with lymphocytes.^37^ The LD1 and LD2 loadings in Fig. 5 share many similarities between OPTIR spectra of cells on CaF_2_ and on glass with 532 nm probe laser which explain the balanced accuracies above 80%. The standard deviations tend to be higher at the key bands 1639, 2852 and 2923 cm^-1^ in second derivatives for granulocytes on glass (Fig. 5f) which is also evident by the larger spread of the scores in the violin plot (Fig. 5d). Most of the LD1 and L2 loading features are conserved even for cells on glass with 785 nm probe laser. The reduced accuracy of 63% is due to the less pronounced variations at the key bands 1639, 2852 and 2923 cm^-1^ in second derivatives. Furthermore, variations appear for LD1 and LD2 loadings in the wavenumber range 1313 to 1500 cm^-1^ that do not seem to be specific.

The transition from CaF_2_ to glass reduces consumable cost and moves OPTIR onto the same substrate used in routine histopathology, a methodological step toward compatibility with clinical workflows. Although glass absorbs between 900 and 1200 cm^−1^, the bands resolved in the 1313-2975 cm^−1^ range provide sufficient contrast for subtype discrimination. Discriminatory features associated with granulocyte versus agranulocyte separation were preserved across both CaF_2_ and glass, and across 532 nm and 785 nm probe wavelengths. This is consistent with the discriminating signals being largely intrinsic to the cells rather than substrate or probe dependent artefacts, and it is an encouraging indication that these features may transfer between instrument configurations, a prerequisite for potential clinical translation, where substrate and laser specifications can vary between sites. Confirmation on larger, donor-diverse cohorts is required before this transferability can be claimed quantitatively, and the biological assignments of the LD loadings should likewise be read as consistent with, rather than proof of, the underlying biochemistry. Substrate and probe wavelength are not fully orthogonal in the present design as the CaF_2_/785 nm combination was not acquired. The reduced lymphocyte–monocyte resolution on glass at 785 nm cannot be attributed unambiguously to the substrate; a probe-wavelength contribution cannot be excluded. Because OPTIR is non-destructive, spectra can be acquired first and the same cells subsequently stained for morphological validation, extending the approach to scenarios where morphology alone is insufficient.

The fluorescence and the spectra age differently, which shapes how the workflow is timed. The OPTIR spectra held their band positions and relative intensities over 14 days of storage (Fig. S1), so classification is stable and the spectral measurement can be delayed without loss of subtype information. The intrinsic fluorescence used to find leukocytes, by contrast, was weak in fresh cells and grew brighter over the same period (Fig. 1d). For clinical use this is a limitation, since smears are read soon after preparation, when the fluorescence contrast is weakest, so an alternative localization route may be needed for freshly prepared blood smear samples such as discrete wavenumber OPTIR images at 1546 and 2922 cm^-1^.

## Conclusions

This study demonstrates that OPTIR spectroscopy successfully classifies granulocytes, lymphocytes and monocytes in unstained peripheral blood smears using their intrinsic biochemical contrast. Discrete wavenumber imaging at 1546 and 2922 cm^-1^ identified leukocytes in full blood smears. Fluorescence microscopy using intrinsic leukocyte fluorescence constitutes a convenient solution for their localization. The intrinsic fluorescence was weak in fresh smears and increased over 14 days of storage, so fluorescence-guided screening is least favourable when smears are read fresh, whereas the OPTIR spectra stayed stable over the same period on both substrates and can be acquired even after 14 days without loss of subtype information. Single-cell acquisition between inherent fluorescence-guided localization and post-acquisition Giemsa staining makes every spectrum traceable to a cytologically verified cell. PCA-LDA gave balanced accuracies of 96% on CaF_2_ and 80% on glass. The granulocyte–mononuclear differentiation along LD1 was preserved across substrates and probe wavelengths, and is driven by lipid, nucleic-acid and protein secondary-structure content; the lymphocyte–monocyte distinction was achieved on CaF_2_ by LD2 of PCA-LDA. Subtype discrimination on standard histopathology glass is preserved despite loss of the 973–1311 cm^−1^ region, and the non-destructive, fixation-free measurement permits direct correlation with established cytological criteria on the same cells. Future work will cover class-balanced sampling, non-linear classifiers, and larger, donor-stratified patient cohorts including pathological states to enable leave-one-donor out validation and a format assessment of independent-donor generalization. Additionally, OPTIR microscopes using laser scanning instead of scanning by a motorized stage and in wide-field configuration that can enable image acquisition within seconds will be explored. Taken together, the results position this work as a methodological proof of concept for OPTIR-based single-cell screening in whole blood smears, on which future clinically oriented studies with larger and pathological cohorts can build.

## Author contributions

Conceptualization and methodology, C.K. and S.R.; investigation and formal analysis, S.R. and C.R.; resources, S.P. and M.K.; writing—original draft, S.R. and C.K.; writing—review & editing, S.R., S.P., C.R. and C.K.; supervision, C.K. and J.P.; funding acquisition, C.K., M.K. and J.P. All authors have read and agreed to the published version of the manuscript.

## Acknowledgements

This work is supported by the BMBF, funding program Photonics Research Germany (13N15464 and 13N15720) and is integrated into the Leibniz Center for Photonics in Infection Research (LPI). The LPI initiated by Leibniz-IPHT, Leibniz-HKI, UKJ and FSU Jena is part of the BMBF national roadmap for research infrastructures. The OPTIR instrument Mirage-R was funded within the Innovation Center ThIMEDOP (2018 IZN 0003). The Mirage-LS instrument was funded by the Free State of Thuringia and cofinanced by the European Union through the European Regional Development Fund (EFRE, Project 2023 HSB 0025). We thank the Institute of Clinical Transfusion Medicine Jena from Jena University Hospital for providing blood smear samples.

## Supplementary Information

**Figure S1.**
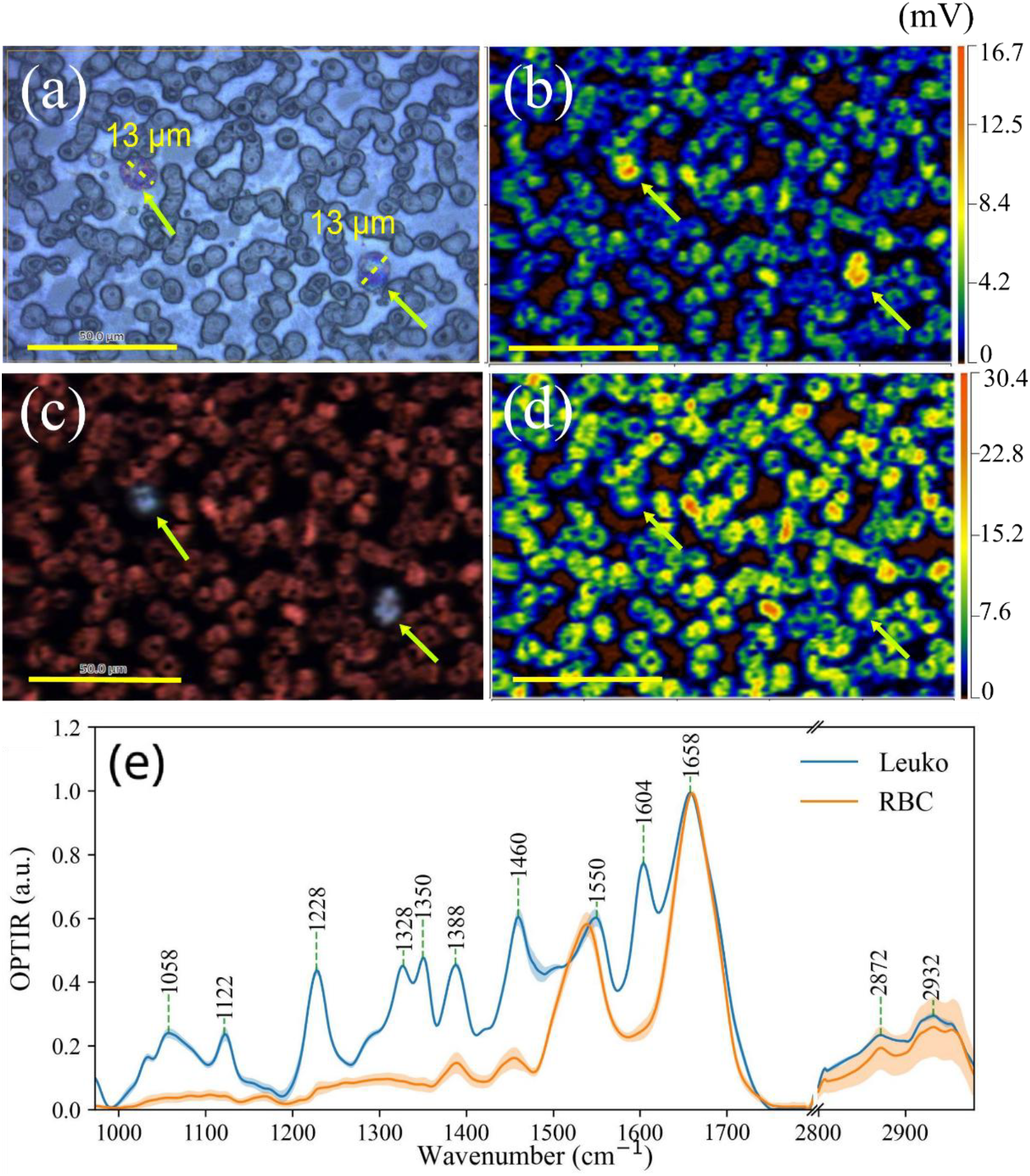
(a) Microscope image of leukocytes and erythrocytes (red blood cells, RBCs) using a 40× objective. (b) OPTIR image of stained blood smears on CaF2 acquired at 1604 cm^-1^. The yellow arrows point to the stained leukocytes. (c) RGB overlay of 1604 cm^-1^ and 1660 cm^-1^ images. (d) OPTIR image acquired at amide I, 1660 cm^-1^. Scale bar in (a) – (d) = 50 µm. (e) Mean OPTIR spectrum of the two leukocytes (blue) highlighted in (a-d) and multiple erythrocytes (RBC, orange) of blood smears on CaF2 after Giemsa staining. Numerous intense bands of the stain are evident in the fingerprint range.

**Figure S2.**
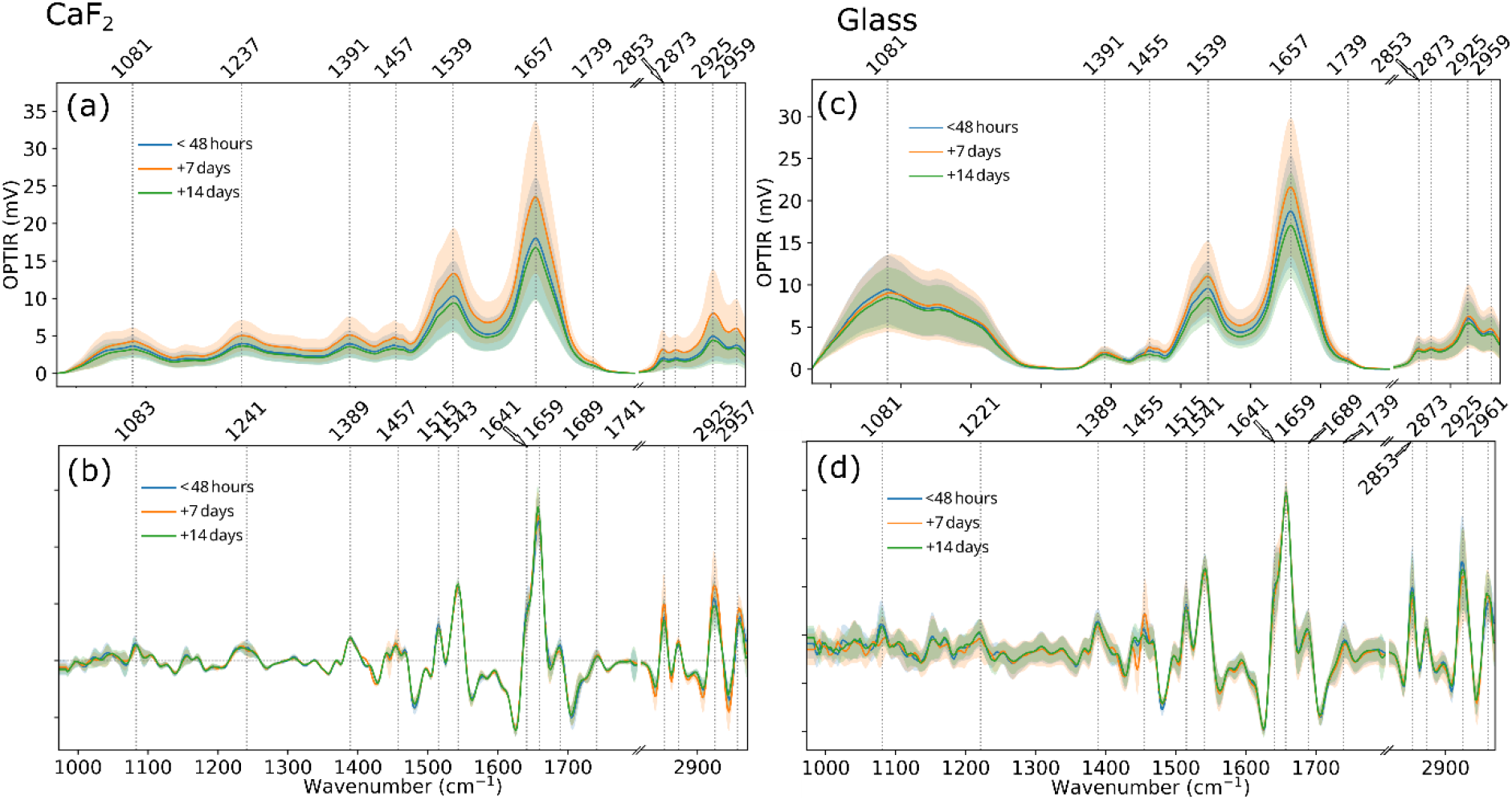
Blood smear time-dependence. (a) CaF2, (c) Glass OPTR spectra at different time points. (b), (d) Second derivative features are well conserved across substrate and time points.

**Figure S3.**
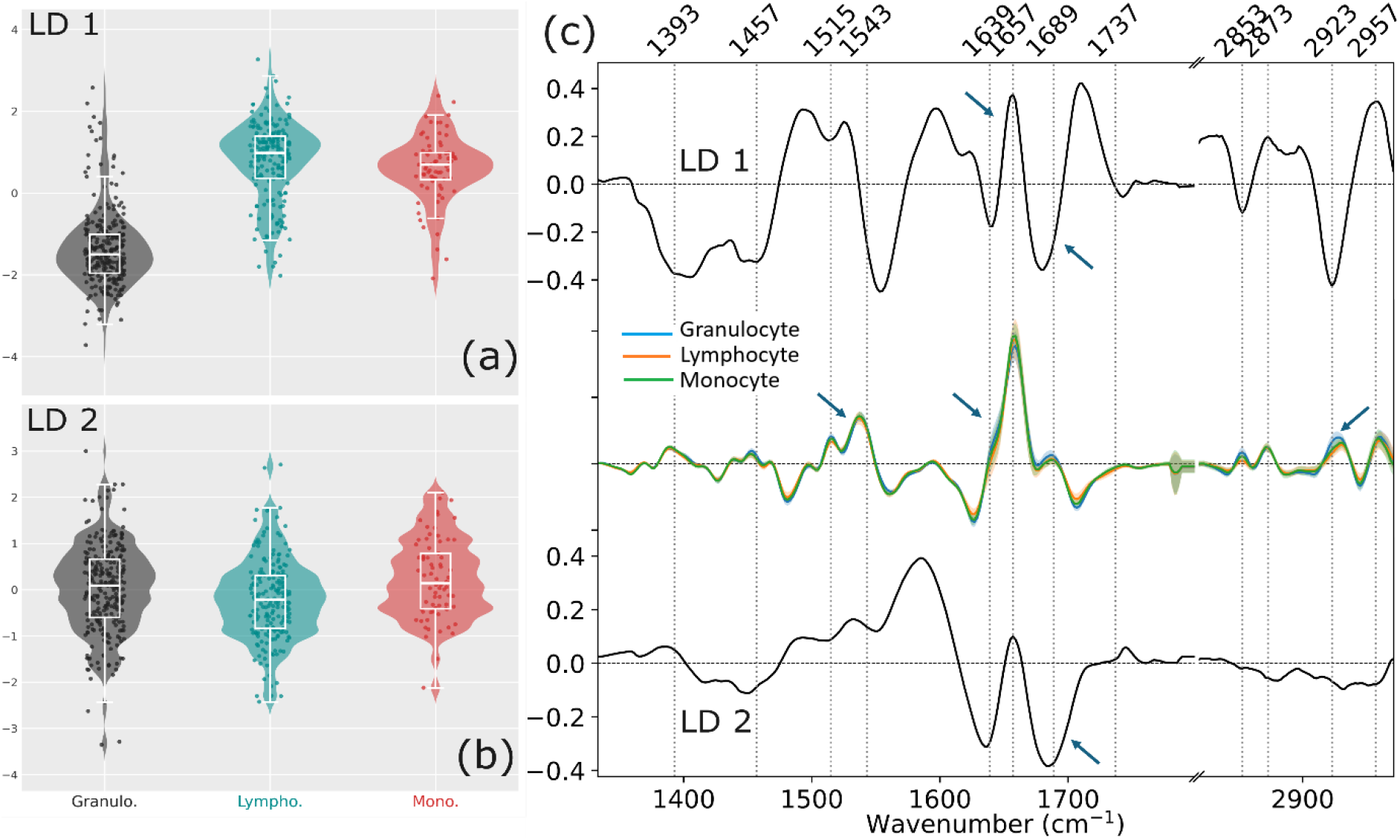
Principal component - linear discriminant analysis (PCA-LDA) analysis of OPTIR spectra from granulocytes (grey), lymphocytes (teal) and monocytes (red measured on glass with a 785 nm probe. (a, b) Violin plots of LD1 and LD2 scores across the three subpopulations. (c) LD1 and LD2 loadings (black) shown alongside the mean negative second-derivative spectra (red) of each leukocyte subpopulation for comparison. Arrows highlight the spectral regions driving class separation.

**Table S1.** Donor, smear, and single cell counts of full blood per acquisition configuration. Cells are per-subtype: G = granulocytes, L = lymphocytes, M = monocytes.

| Configuration | Substrate | Probe (nm) | Donors | Smears | Cells (G / L / M) | Total cells |
| --- | --- | --- | --- | --- | --- | --- |
| CaF<sub>2</sub>/532 | CaF<sub>2</sub> | 532 | 1 | 1 | 50 / 30 / 20 | 100 |
| Glass/532 | glass | 532 | 1 | 2 | 100 / 60 / 40 | 200 |
| Glass/785 | glass | 785 | 7 | 7 | 260 / 203 / 73 | 536 |

**Table S2.** OPTIR band assignments. Leukocyte band assignments in mean OPTIR spectra, second derivatives (polarity reversed) and linear discriminants^1,2^.

| LD 1 / LD 2 (cm<sup>-1</sup>) | Second Derivative (cm<sup>-1</sup>) | Assignment |
| --- | --- | --- |
| **1391/1393** | 1389 | $\nu_s$ (COO<sup>-</sup>)/ $\delta$(CH<sub>2</sub>); Proteins |
| **1457** | 1457 | $\delta$(CH<sub>2</sub>)/ $\delta$(CH<sub>2</sub>,CH<sub>3</sub>); Proteins/Lipids |
| **1513, 1531** | 1543 | amide II ($\delta$(N-H) + $\nu$(C-N)); Proteins |
| **1639** | 1641 | amide I, β-sheet; Proteins (most prominent in the Granulocytes) |
| **1657** | 1657 | amide I, α-helix; Proteins |
| **1688** | 1689 | $\nu$(C=O); Nucleic acid base |
| **1737** | 1741/1743 | $\nu$(C=O); Phospholipids |
| **2853** | 2853 | $\nu_s$ (CH<sub>2</sub>); Lipids |
| **2873** | 2873 | $\nu_s$ (CH<sub>3</sub>); Lipids |
| **2923** | 2923 | $\nu_{as}$ (CH<sub>2</sub>); Lipids |
| **2957** | 2957 | $\nu_{as}$ (CH<sub>3</sub>); Lipids |

